# Skipping a beat: heartbeat-evoked potentials reflect predictions during interoceptive-exteroceptive integration

**DOI:** 10.1101/2020.04.21.053173

**Authors:** Leah Banellis, Damian Cruse

**Affiliations:** School of Psychology and Centre for Human Brain Health, University of Birmingham, Edgbaston, B15 2TT, UK

**Keywords:** Attention, expectation, interoception, predictive coding, precision

## Abstract

Several theories propose that emotions and self-awareness arise from the integration of internal and external signals and their respective precision-weighted expectations. Supporting these mechanisms, research indicates that the brain uses temporal cues from cardiac signals to predict auditory stimuli, and that these predictions and their prediction errors can be observed in the scalp heartbeat-evoked potential (HEP). We investigated the effect of precision modulations on these cross-modal predictive mechanisms, via attention and interoceptive ability. We presented auditory sequences at short (perceived synchronous) or long (perceived asynchronous) cardio-audio delays, with half of the trials including an omission. Participants attended to the cardio-audio synchronicity of the tones (internal attention) or the auditory stimuli alone (external attention). Comparing HEPs during omissions allowed for the observation of pure predictive signals, without contaminating auditory input. We observed an early effect of cardio-audio delay, reflecting a difference in heartbeat-driven expectations. We also observed a larger positivity to omissions of sounds perceived as synchronous than to omissions of sounds perceived as asynchronous when attending internally only, consistent with the role of attentional precision for enhancing predictions. These results provide support for attentionally-modulated cross-modal predictive coding, and suggest a potential tool for investigating its role in emotion and self-awareness.

---

The Bayesian brain hypothesis states that the brain is a probabilistic machine, with hierarchical neuronal representations underlying cognition, perception, and behaviour (Friston, 2009). The predictive coding framework posits that, in the comparison between top-down predictions from high-level brain regions and incoming low-level sensory input, any difference between the two signals is propagated up the hierarchy as a prediction error, thus allowing for iterative updating of the higher-level representations (Rao & Ballard, 1999). Successful matching of predictions with incoming stimuli, and thus successful minimisation of prediction error, results in ‘correct’ perception, cognition, and action (Friston, 2010). Minimisation of prediction error is accomplished either by updating predictive models to accommodate unexpected signals (i.e. perceptual inference) or by performing actions (such as motor or autonomic responses) to better match predictions (i.e. active inference) (Adams, Shipp, & Friston, 2013; Friston, 2010), consistent with an embodied view of cognition (Allen & Friston, 2018).

As with perception of external stimuli (exteroception), perception of internal stimuli (interoception) is also considered to be supported by hierarchical prediction error minimisation mechanisms (Barrett & Simmons, 2015; Seth, 2013; Seth, Suzuki, & Critchley, 2012). Broadly, interoception is the perception of visceral bodily sensations such as heartbeat contractions, the expansion of lungs, or feelings of the body’s internal state such as hunger or nausea (Cameron, 2001; Sherrington, 1948). The Embodied Predictive Interoceptive Coding (EPIC) model describes an interoceptive cortical network comprising of viscerosensory neural afferents which arrive at the brainstem and thalamus via the dorsal root ganglion and vagus nerve, outputting to the hypothalamus, amygdala, anterior cingulate cortex and the insula, with its highest regions residing in the posterior ventral medial prefrontal cortex (vmPFC) and the orbitofrontal cortex (Barrett & Simmons, 2015; Critchley & Harrison, 2013; Damasio & Carvalho, 2013; Quadt, Critchley, & Garfinkel, 2018). This network is thought to be involved in numerous high level cognitive processes such as emotional processing, bodily self-consciousness, visual awareness, self-recognition, attention and time perception (Azzalini, Rebollo, & Tallon-Baudry, 2019; Craig, 2009; Critchley & Harrison, 2013; Quadt et al., 2018; Tsakiris & Critchley, 2016). Indeed, as part of a prediction error minimisation framework, Seth et al (2012; 2013) have proposed that embodied selfhood and emotional experience are the outcome of successful suppression of interoceptive prediction errors through active inference (Seth & Friston, 2016). Additionally, dysfunctional interoceptive predictive mechanisms have been proposed to account for a variety of psychological disorders such as anxiety, depression, autism, dissociative disorders, and psychotic illnesses (Haker, Schneebeli, & Stephan, 2016; Quattrocki & Friston, 2014; Seth & Friston, 2016; Seth et al., 2012), thus increasing scientific interest in characterising these mechanisms.

One potential method of investigating the neural basis of interoceptive predictive mechanisms is by analysing heart-evoked potentials (HEPs) (Pollatos & Schandry, 2004; Schandry, Sparrer, & Weitkunat, 1986). HEPs are averaged electrophysiological signals time-locked to heartbeats and are thought to reflect neuronal processing of cardiac afferents, encompassing interoceptive prediction error of each individual heartbeat (Ainley, Apps, Fotopoulou, & Tsakiris, 2016; Petzschner et al., 2019). In a recent study on interoceptive predictions, Pfeiffer & De Lucia (2017) presented healthy participants with a sequence of tones that were either synchronous or asynchronous with their own heartbeat. Crucially, the occasional tone was unexpectedly omitted from these sequences. Evoked responses to expected sounds that did not happen – i.e. omission responses – are an elegant way of observing pure prediction signals without the contamination of auditory potentials (Chennu et al., 2016; Wacongne et al., 2011). Consequently, Pfeiffer & De Lucia (2017) reported a larger HEP during omission periods in cardiac synchrony, relative to cardiac asynchrony, consistent with a predictive account in which the brain uses the interoceptive (cardiac) signals to predict upcoming exteroceptive signals (sounds).

Predictions and their errors are also influenced by their precision – formally, the inverse of the variance, or the uncertainty in the signal. Within the prediction error minimisation framework, attention is described as a means to optimise the relative precision weight of predictions and prediction error signals, via synaptic gain control (Friston, 2009). For example, attending to a specific sensory signal is thought to enhance the precision of the predictions related to that signal, subsequently influencing associated prediction errors (Hohwy, 2012). Consistent with the characterisation of the HEP as a neural correlate of precision-weighted interoceptive prediction error, many studies have reported attentional modulation of the amplitude of the HEP – for example, during tasks involving attending to heartbeat sensations relative to external stimuli (García-Cordero et al., 2017; Montoya, Schandry, & Müller, 1993; Petzschner et al., 2019; Schandry et al., 1986; Villena-González et al., 2017; Yuan, Yan, Xu, Han, & Yan, 2007).

The relative weight of precision in perceptual inference is also influenced by individual differences in relative uncertainty (Lawson, Rees, & Friston, 2014; Seth & Friston, 2016). For example, individuals who are accurate at identifying when sounds are synchronous with their heartbeat (i.e. performance on the heartbeat detection task) also exhibit higher HEP amplitudes relative to individuals who are poor heartbeat perceivers, just as in an attentive versus inattentive contrast (Katkin, Cestaro, & Weitkunat, 1991; Pollatos, Kirsch, & Schandry, 2005; Pollatos & Schandry, 2004; Schandry et al., 1986). Indeed, Ainsley et al. (2016) have previously characterised these individual differences in interoceptive ability as individual differences in relative precision of prediction errors. However, caution should be taken when interpreting differences across interoceptive ability groups, as multiple heartbeat detection paradigms exist, which assess distinct processes and may not measure interoceptive ability validly (Brener & Ring, 2016; Corneille et al., 2020; Ring & Brener, 2018). In addition, Garfinkel et al. (2015) suggested three distinct and dissociable dimensions of interoceptive ability: interoceptive sensibility, accuracy, and awareness, with each dimension potentially influencing predictive mechanisms differently.

Consequently, a combined study of attention to interoceptive signals and individual differences in interoceptive ability allows us to directly test this predictive framework within the domain of evoked potentials. Specifically, here we report the effect of attention and interoceptive ability on interoceptive predictions reflected in the electrical potentials evoked by omissions within a heartbeat detection task. As omission-evoked responses reflect top-down predictions from higher cortical regions, our approach allows us to measure the influence of attentional precision on interoceptive prediction and error signals, without contaminating bottom-up input (Chennu et al., 2016; Wacongne et al., 2011). Consistent with characterisations of the precision-weighting nature of both within-subject and between-subject variations in attention (Chennu et al., 2016; Feldman & Friston, 2010; Hohwy, 2012), we hypothesised that HEPs during auditory omission periods would be 1) larger when sounds are perceived as synchronous with the heartbeat, 2) larger when the heartbeat is attended, and 3) larger for those individuals with high interoceptive ability. At the source level, we anticipated increased anterior insula activation when sounds are perceived as synchronous, supporting the role of the insula as a hub for interoceptive and exteroceptive integration (Gray, Harrison, Wiens, & Critchley, 2007; Salomon et al., 2016). Furthermore, we hypothesised increased activation in the insula, cingulate cortex, and somatosensory cortex (postcentral gyrus) when directing attention internally, than externally, and in individuals with high interoceptive perception, than poor interoceptive perceivers, as previously observed in fMRI studies (Critchley, Wiens, Rotshtein, Öhman, & Dolan, 2004; García-Cordero et al., 2017).

## Materials and Methods

Unless otherwise stated, all methods, analyses, and hypotheses were pre-registered at [https://osf.io/nr8my/].

### Participants

We recruited 39 participants from the University of Birmingham via advertisement on posters or the online SONA Research Participation Scheme. Our inclusion criteria were: righthanded 18 to 35-year-olds, with no reported cardiovascular or neurological disorders. We compensated participants with course credit. The Psychology Research Ethics Board of the University of Birmingham granted ethical approval for this study and written informed consent was completed by all participants. The data of five participants were excluded because of poor data quality resulting in more than a third of the trials of interest rejected. Subsequent analyses were completed on a final sample of 34 participants (Median age = 20 years, Range = 18-28 years). We chose this sample size in advance as it provides 80% power to detect a medium effect size (0.5) in our within-subjects interaction between attention and cardio-audio delay (alpha=.05; GPower, Faul et al., 2007).

### Stimuli and Procedure

The experiment consisted of four blocks of 56 trials (224 trials total), with each trial consisting of 7 to 10 auditory tones (1000Hz, 100ms duration, 44100 sampling rate) presented via external speakers, with breaks given between each block. The onset of each tone was triggered by the online detection of the participants R-peak from electrocardiography (ECG) recordings using Lab Streaming Layer and a custom MATLAB script (Kothe et al., 2018). The script analysed in real time the raw ECG signal by computing the variance over the preceding 33ms window and determining if the signal exceeded an individually adjusted threshold, at which point a tone was triggered to occur after either a 250ms (perceived synchronous) or 550ms (perceived asynchronous) delay (Brener & Kluvitse, 1988; Wiens & Palmer, 2001). In half of the trials, the third from last tone was omitted, resulting in an R-peak without an auditory stimulus. A fixation cross was present during tone presentation.

A cue at the start of each trial (200ms) directed participants’ attention to focus internally (‘Heart’) or externally (‘Tone’). During the internal task, participants focused on their heartbeat sensations (without taking their pulse), and determined whether the tones presented were synchronous or not with their heartbeat. During the external task, participants were told to ignore their heartbeat sensations and direct attention towards the sounds alone. The external task was to determine whether there was a missing sound during that trial. Participants responded to the internal task (‘Were The Tones Synchronous With Your Heart?’) or external task (‘Was There a Missing Tone?’) question by pressing ‘y’ for yes or ‘n’ for no on the keyboard, and rated their confidence in their decision from 1 to 4 (1 = total guess, 2 = somewhat confident, 3 = Fairly Confident, 4 = Complete Confidence). The inter-trial interval was between 2 to 3 seconds, chosen from a uniform distribution on each trial (see Figure 1). The order of the experimental conditions were randomized to ensure no more than 3 of the same condition on consecutive trials. Finally, participants completed the short Porges Body Perception Questionnaire (BPQ), including a body awareness and autonomic reactivity subscale (Porges, 1993).

**Figure 1.**
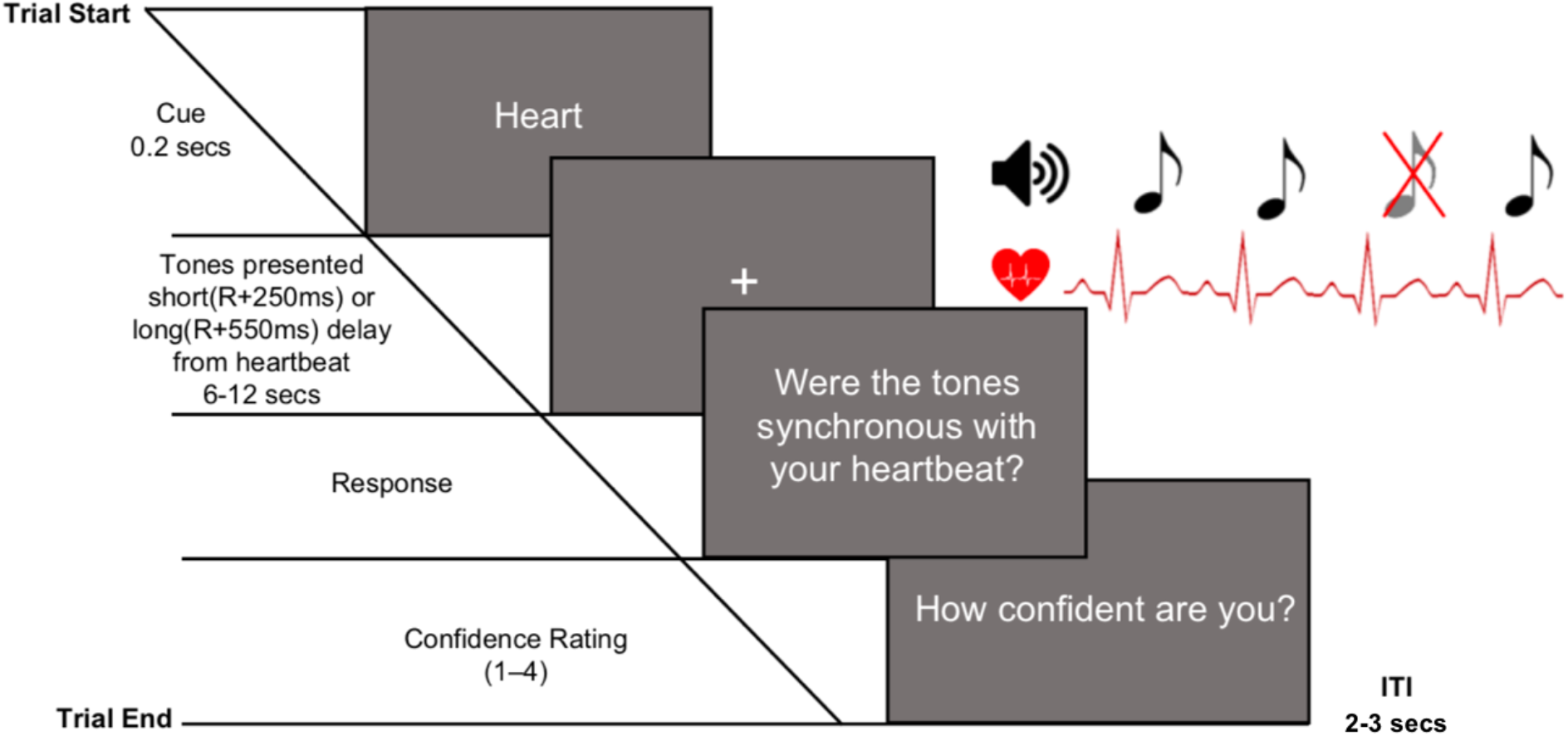
Experimental design of the integrated heartbeat detection (internal attention) and omission detection task (external attention), displaying an internal attention trial.

### Indices

Interoceptive accuracy was calculated by comparing the normalised proportion of hits (responding ‘yes’ to a short cardio-audio delay ‘R+250ms’ trial) with the normalised proportion of false alarms (responding ‘yes’ to a long cardio-audio delay ‘R+550ms’ trial) (i.e. d-prime *(d)*) (Macmillan & Creelman, 1990). The proportion of hits and false alarms were normalised using the inverse of the standard normal cumulative distribution. As in previous studies (Ewing et al., 2017; Garfinkel et al., 2015), we quantified sensibility to a variety of internal bodily sensations with the score on the awareness subsection of the Porges Body Perception Questionnaire (BPQ) (Porges, 1993) and defined sensibility to heartbeat sensations as the median confidence rating during internal trials (Ewing et al., 2017; Forkmann et al., 2016; Garfinkel et al., 2015).

Interoceptive awareness was calculated using type 2 signal detection theory analysis comparing observed type 2 sensitivity (meta-*d*’) with expected type 2 sensitivity (*d*’) (Maniscalco & Lau, 2012). Meta-*d*’ is the *d*’ expected to generate the observed type 2 hit rates and type 2 false alarm rates and was estimated using maximum likelihood estimation (MLE) (Maniscalco & Lau, 2014). This determined the extent to which confidence ratings predicted heartbeat detection accuracy, and thus interoceptive awareness. Groups were separated into high/low interoceptive accuracy, sensibility, and awareness with median splits.

### EEG/ECG acquisition

EEG was recorded throughout the experiment using a gel-based 128-channel Biosemi ActiveTwo system, acquired at 512Hz, referenced to the Common Mode Sense electrode located approximately 2-cm to the left of CPz. Two additional electrodes recorded data from the mastoids, and ECG was measured using two electrodes placed on either side of the chest, also sampled at 512Hz.

### EEG/ECG Pre-Processing

First, we filtered the continuous EEG data in two steps (i.e. high-pass then low-pass) between 0.5Hz and 40Hz using the finite impulse response filter implemented in EEGLAB (function: pop_eegfiltnew). We filtered ECG between 0.5Hz and 150Hz (Kligfield et al., 2007). Next, we segmented the filtered EEG signals into epochs from −300ms to 800ms relative to the R-peak of the ECG recording during the omission period, re-referenced to the average of the mastoids. We detected the R-peaks using a custom MATLAB script, and subsequently checked the accuracy of R-peak detection via visual inspection. When necessary, we manually corrected the estimated R-peaks to ensure accurate R-peak detection. Any trials with missed or multiple sounds per R-peak were rejected. The subsequent artefact rejection proceeded in the following steps based on a combination of methods described by Nolan et al., 2010 and Mognon et al., 2011 (Mognon, Jovicich, Bruzzone, & Buiatti, 2011; Nolan, Whelan, & Reilly, 2010).

First, bad channels were identified and removed from the data. We consider a channel to be bad if its absolute z-score across channels exceeds 3 on any of the following metrics: 1) variance of the EEG signal across all time-points, 2) mean of the correlations between the channel in question and all other channels, and 3) the Hurst exponent of the EEG signal (estimated with the discrete second order derivative from the Matlab function *wfbmesti).* After removal of bad channels, we identified and removed trials containing non-stationary artefacts. Specifically, we considered a trial to be bad if its absolute z-score across trials exceeds 3 on any of the following metrics: 1) the mean across channels of the voltage range within the trial, 2) the mean across channels of the variance of the voltages within the trial, and 3) the mean across channels of the difference between the mean voltage at that channel in the trial in question and the mean voltage at that channel across all trials. After removal of these individual trials, we conducted an additional check for bad channels, and removed them, by interrogating the average of the channels across all trials (i.e. the ERP, averaged across all conditions). Specifically, we considered a channel to be bad in this step if its absolute z-score across channels exceeds 3 on any of the following metrics: 1) the variance of voltages across time within the ERP, 2) the median gradient of the signal across time within the ERP, and 3) the range of voltages across time within the ERP.

To remove stationary artefacts, such as blinks and eye-movements, the pruned EEG data is subjected to independent component analysis with the *runica* function of EEGLAB. The Matlab toolbox ADJUST subsequently identified which components reflect artefacts on the basis of their exhibiting the stereotypical spatio-temporal patterns associated with blinks, eye-movements, and data discontinuities, and the contribution of these artefact components is then subtracted from the data (Mognon et al., 2011). Next, we interpolated the data of any previously removed channels via the spherical interpolation method of EEGLAB, and rereferenced the data to the average of the whole head.

We included an additional preprocessing step beyond those planned in our preregistration to control for differences in the cardiac field artefact (CFA) at our different delay conditions (Nakamura & Shibasaki, 1987). Specifically, we calculated single-subject average HEPs during rest periods, following the same preprocessing pipeline as the experimental HEPs. In a similar approach to that used in previous research (Van Elk et al., 2014), we then subtracted the average resting HEP from individual experimental trials, locked to each heartbeat. This conservative method eliminates remaining artefacts due to additional heartbeats within the same trial.

Before proceeding to group-level analyses, single-subject CFA-corrected averages for HEP analysis are finalised in the following way. First, a robust average was generated for each condition separately, using the default parameters of SPM12. Robust averaging iteratively down-weights outlier values by time-point to improve estimation of the mean across trials. As recommended by SPM12, the resulting HEP was low-pass filtered below 20Hz (again, with EEGLAB’s pop_neweegfilt), and the mean of the baseline window (−100ms – 0ms) was subtracted.

### HEP Analysis

HEPs during the omission period were compared with the cluster mass method of the open-source Matlab toolbox FieldTrip (Oostenveld, Fries, Maris, & Schoffelen, 2011: fieldtrip-20181023). This procedure involves an initial parametric step followed by a non-parametric control of multiple-comparisons (Maris & Oostenveld, 2007). Specifically, we conducted either two-tailed dependent samples t-tests (for comparison 1) or a combination of two-tailed independent and dependent samples t-tests (for comparison 2) at each spatio-temporal data-point within the time window. Spatiotemporally adjacent t-values with p-values < 0.05 are then clustered based on their proximity, with the requirement that a cluster must span more than one time-point and at least 4 neighbouring electrodes, with an electrode’s neighbourhood containing all electrodes within a distance of .15 within the Fieldtrip layout coordinates (median number of neighbours = 11, range 2-16). Finally, we summed the t-values at each spatio-temporal point within each cluster. Next, we estimated the probability under the null hypothesis of observing cluster sum Ts more extreme than those in the experimental data – i.e. the p-value of each cluster. Specifically, Fieldtrip randomly shuffles the trial labels between conditions, performs the above spatio-temporal clustering procedure, and retains the largest cluster sum T. Consequently, the p-value of each cluster observed in the data is the proportion of the largest clusters observed across 1000 such randomisations that contain larger cluster sum T’s.

Our pre-registered analyses were to be conducted on the ERP data from 100ms to 600ms relative to the R-peak. However, it subsequently became evident that this approach is confounded by the lag difference in tone presentation across conditions. Consequently, here we report one set of analyses on ERP data from 0ms to 250ms post-R (i.e. before any anticipated tone would have occurred) and a second set of analyses from 0ms to 250ms relative to the onset of the omitted sound (i.e. from 250ms to 500ms post-R for the short delay condition, and 550ms to 800ms post-R for the long delay condition).

### Comparisons

Using the above method, HEPs were compared across cardio-audio delay and attention conditions to assess the main effects, and the interaction was calculated as the difference between short-delay and long-delay trials between attention groups (comparison 1). If an interaction was observed, pairwise separate analyses were completed to consider simple effects. Similar comparisons were completed across attention and interoceptive individual difference conditions (interoceptive awareness, accuracy and sensibility) (comparison 2).

### CFA Control Analyses

We performed control analyses on the ECG data, to determine if differences in cardiac activity contributed towards the HEP results. Therefore, equivalent analyses to that performed on the HEPs were completed on the ECG data. Subsequently, single-subject robust averages of the ECG activity were computed for each condition and were analysed using the cluster mass method, as described above. The same comparisons were completed as to those which showed a significant HEP effect (i.e. ECG was compared across cardio-audio delay conditions 0-250ms post-R, and the attention and delay interaction was assessed 0-250ms relative to the omission).

### Source Reconstruction

Since our initial pre-registration, we discovered that our planned source analysis pipeline performed poorly at localising basic sensory responses in a separate study in our lab. Consequently, we concluded that those pre-registered methods were inappropriate for this study. Therefore, here we report a more rudimentary but validated source reconstruction method, using statistical parametric mapping (SPM12) (Henson et al., 2009; López et al., 2014).

Our source estimation approach was completed for each time-window separately in which we observed a significant sensor level effect: 19-251ms post-R for the main effect of delay and 136-177ms relative to the omission for the attention and delay interaction (i.e. 386-427ms post-R for the short delay condition and 686-727ms post-R for the long delay condition). For each time-window, within SPM12, we applied a hanning taper to downweight the signal at the beginning and end of the window in the condition-wise grand averages, and filtered the data between 1 and 48 Hz. Cortical sources of each sensor-level HEP were reconstructed using the default anatomical template in SPM. Electrode positions were coregistered to the template using the fiducials of the nasion, left peri-auricular and right periauricular points. We calculated the forward model using the Boundary Element Method. The inverse model was generated based on an empirical Bayesian approach. Specifically, we applied the greedy search fitting algorithm, which optimises the multiple sparse priors approach when localising the sensor-level evoked responses. Finally, we contrasted the condition-wise source estimates (i.e. generated difference source volumes). The estimated source results were projected onto a canonical inflated brain surface for visualisation, using the open source MNI2FS toolbox (Price., 2020: https://github.com/dprice80/mni2fs).

## Results

### Behavioural data

Participants’ interoceptive accuracy scores (*d*’) were significantly greater than zero (M=0.218, SD=0.347; t(33) = 3.665, *p* < .001). This indicates that performance on the heartbeat discrimination task was above chance and therefore confirms our interpretation of the R+250ms cardio-audio delay as perceived synchronous and R+550ms as perceived asynchronous.

### Heart-evoked potentials Cardio-audio expectation

We observed a significant early dipolar main effect of cardio-audio delay (positive cluster *p* = .002, and negative cluster *p* = .007), perhaps reflecting a difference in expectation induced by the heartbeat. Estimated generators of this effect include bilateral primary somatosensory cortex, bilateral primary motor cortex, right anterior prefrontal cortex, and left ventral temporal cortex. The positive cluster extended from 19-251ms and the negativity cluster 89-251 ms post R-peak, reflecting positive-going waveforms in the short delay condition and negative-going waveforms in the long delay condition. We observed no significant main effect of attention on pre-omission responses (smallest cluster *p* = .094) (see Figure 2).

**Figure 2:**
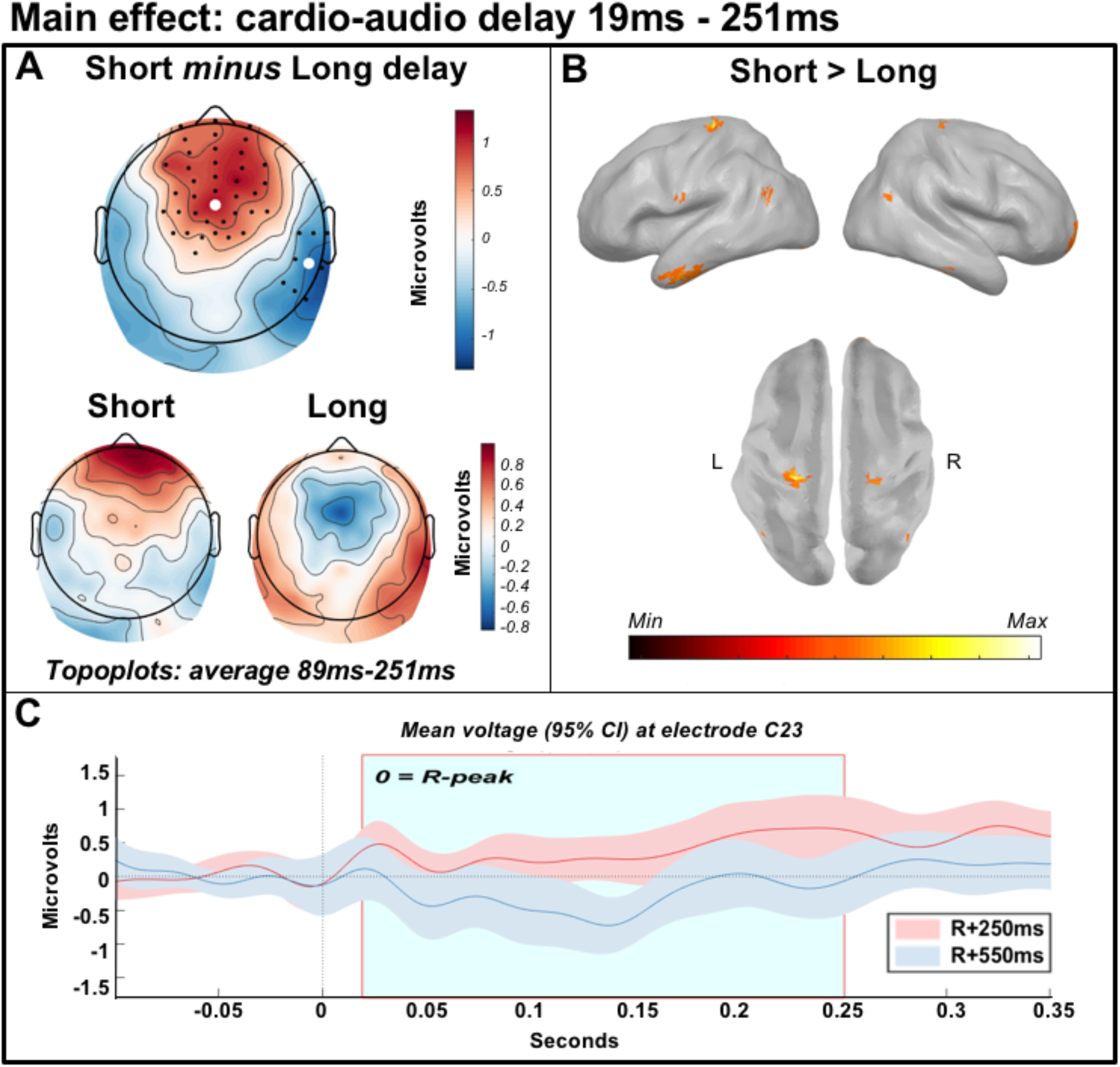
Main effect of cardio-audio delay from 19ms-251ms, reflecting differences in cardio-audio expectation. [A] Scalp distribution of the average significant difference across delay conditions 89-251ms, with electrodes contributing to the dipolar clusters marked. [B] Estimated sources of the main effect in bilateral primary somatosensory cortex, bilateral primary motor cortex, right anterior prefrontal cortex and left ventral temporal cortex. [C] Average HEP across participants at electrode C23, light blue shaded region represents the time of the significant positive effect.

### Unfulfilled expectation

The cluster-based permutation test indicated a significant, though weak, interaction between cardio-audio delay and attention (cluster *p* = .017) with estimated sources in right inferior frontal gyrus, right anterior prefrontal cortex, bilateral primary motor cortex, bilateral premotor cortex, left extrastriate cortex and left ventral temporal cortex. The cluster in the observed data extended from 136-177ms post omission. Follow-up simple effects tests indicated a larger positivity within this cluster for short-delay omissions relative to long-delay omissions during internal attention only (*p* = .006), while there were no clusters formed when contrasting the cardio-audio delay conditions when externally attending. Source analyses estimated internal simple effects in bilateral primary motor cortex, bilateral premotor cortex, right inferior frontal gyrus and bilateral anterior prefrontal cortex, left ventral temporal cortex and left extrastriate cortex, while external simple effects were estimated in bilateral premotor cortex, right inferior frontal gyrus and bilateral anterior prefrontal cortex, left extrastriate cortex, right visual association area (see Figure 3).

**Figure 3:**
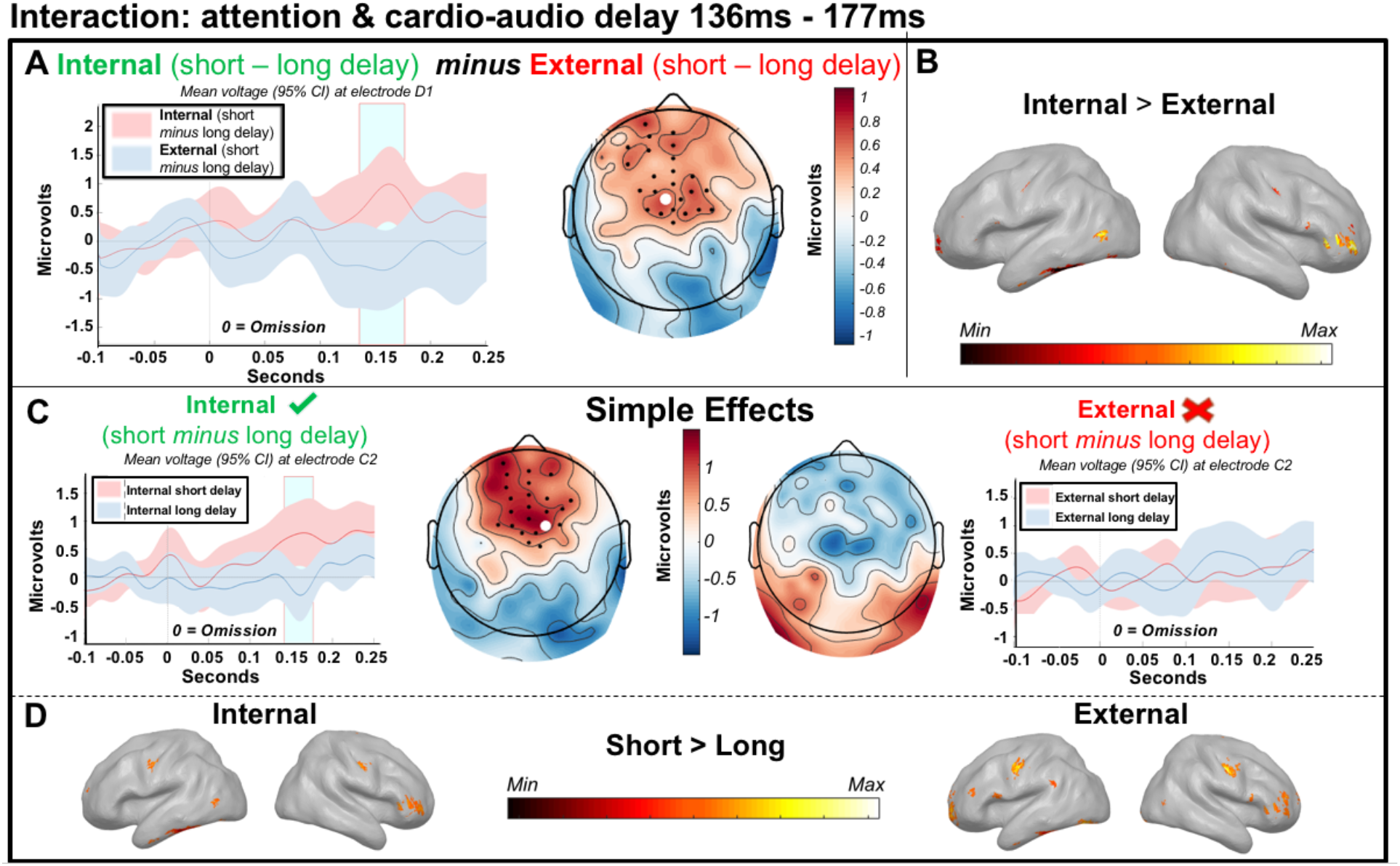
Interaction between attention and cardio-audio delay from 136ms-177ms, relative to the omitted sound. [A - left] Average omission evoked response across participants at electrode D1, light blue shaded region represents the time of the significant effect. [A - right] Scalp distribution of the average significant interaction (attention x delay) 136ms-177ms, with electrodes contributing to the cluster marked. [B] Estimated sources of the interaction in the right inferior frontal gyrus, right anterior prefrontal cortex, bilateral primary motor cortex, bilateral premotor cortex, left extrastriate cortex and left ventral temporal cortex. [C] Analysis of the simple effects showing qualitatively different topographical distributions across attention conditions, and a significant effect of delay in the internal attention condition only. [D - left] Estimated sources of internal simple effects analysis in bilateral primary motor cortex, bilateral premotor cortex, right inferior frontal gyrus and bilateral anterior prefrontal cortex, left ventral temporal cortex and left extrastriate cortex. [D - right] Estimated sources of external simple effects analysis in bilateral premotor cortex, right inferior frontal gyrus and bilateral anterior prefrontal cortex, left extrastriate cortex, right visual association area.

### Control ECG comparisons

We observed no difference clusters when comparing ECG responses between cardio-audio delay conditions, 0-250ms post-R. Similarly, no clusters were found when analysing the interaction between attention and cardio-audio delay on ECG responses, 0-250ms relative to the omitted sound. Therefore, we conclude that it is unlikely that ECG activity contributed towards the HEP differences observed.

### Interoceptive ability

Cluster-based permutation tests indicated no significant interaction of high and low interoceptive awareness (smallest *p* = .128), accuracy (smallest *p* = .043) or sensibility (both median confidence rating and the awareness subsection of the BPQ; smallest *p* = .318) with attention, during short delay trials. Similarly, we observed no significant main effects of interoceptive awareness, accuracy, or sensibility during short delay trials.

We also completed exploratory correlations of interoceptive ability with the amplitude of each participant’s delay effect during the interaction time window (136ms-177ms). These analyses reveal a significant correlation between the delay effect of each participant and interoceptive accuracy during external attention (*r*(32) = −.439, *p* = .009), however this fails to pass a Bonferroni-corrected significance threshold (corrected alpha = .006). We observe no significant correlation between the delay effect and interoceptive awareness during external attention (*r*(32) = .305, *p* = .079), or the delay effect and interoceptive accuracy (*r*(32) = −.193, *p* = .275) or awareness (*r*(32) = −.005, *p* = .980) during internal attention. Additionally, no significant correlations were found with interoceptive sensibility (both the awareness subsection score and the autonomic reactivity subsection score of the BPQ) (smallest *p* = .157) for both internal and external trials (see Figure 4).

**Figure 4:**
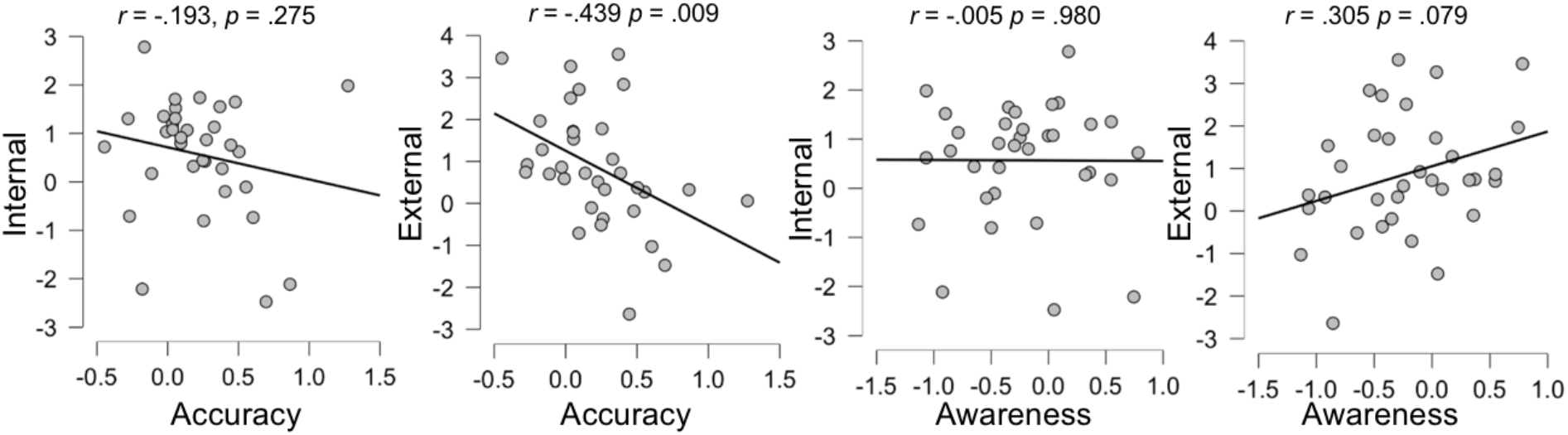
Correlations of interoceptive accuracy and interoceptive awareness with the mean difference in voltage across cardio-audio delay conditions during the significant interaction window for internal and external trials.

### Interbeat intervals (IBIs)

Because previous research found differences in the interbeat intervals following omissions and deviant stimuli, we additionally investigated this as an exploratory analysis (Pfeiffer & De Lucia, 2017; Raimondo et al., 2017). IBIs were significantly longer during internal attention (M=834.356ms, SD=108.000ms) than during external attention (M=813.442ms, SD=102.074ms; F(1,33) = 69.475, *p* <.001, partial n^2^ = .678). However, there was no significant IBI difference between the cardio-audio delay conditions (F(1,33) = 2.342, *p* =.135, partial n^2^ =.066), nor was there a significant interaction between attention and cardio-audio delay (F(1,33) = 3.223, *p* =.082, partial n^2^ =.089).

Additionally, we calculated the IBI’s relative to the omission, revealing an IBI increase post-omission when attending externally. A three-way ANOVA analysed the IBIs post-omission (IBI ‘omission to 1 ’ and IBI ‘1 to 2’) across cardio-audio delay and attention conditions (see Figure 5). This revealed a main effect of IBI (F(1,33) = 17.320, *p* <.001, partial n^2^ = .344), a main effect of attention (F(1,33) = 25.391, *p* <.001, partial n^2^ = .435), a main effect of delay (F(1,33) = 4.605, *p* =.039, partial n^2^ = .122), a significant synchrony and attention interaction (F(1,33) = 5.062, *p* =.031, partial n^2^ = .133) and a significant attention and IBI interaction (F(1,33) = 13.717, *p* <.001, partial n^2^ = .294). The synchrony and IBI interaction was not significant (F(1,33) = 0.339, *p* =.565, partial n^2^ = .010), and the synchrony, attention and IBI interaction was not significant (F(1,33) = 0.007, *p* =.932, partial n^2^ <.001) (see Figure 5).

**Figure 5:**
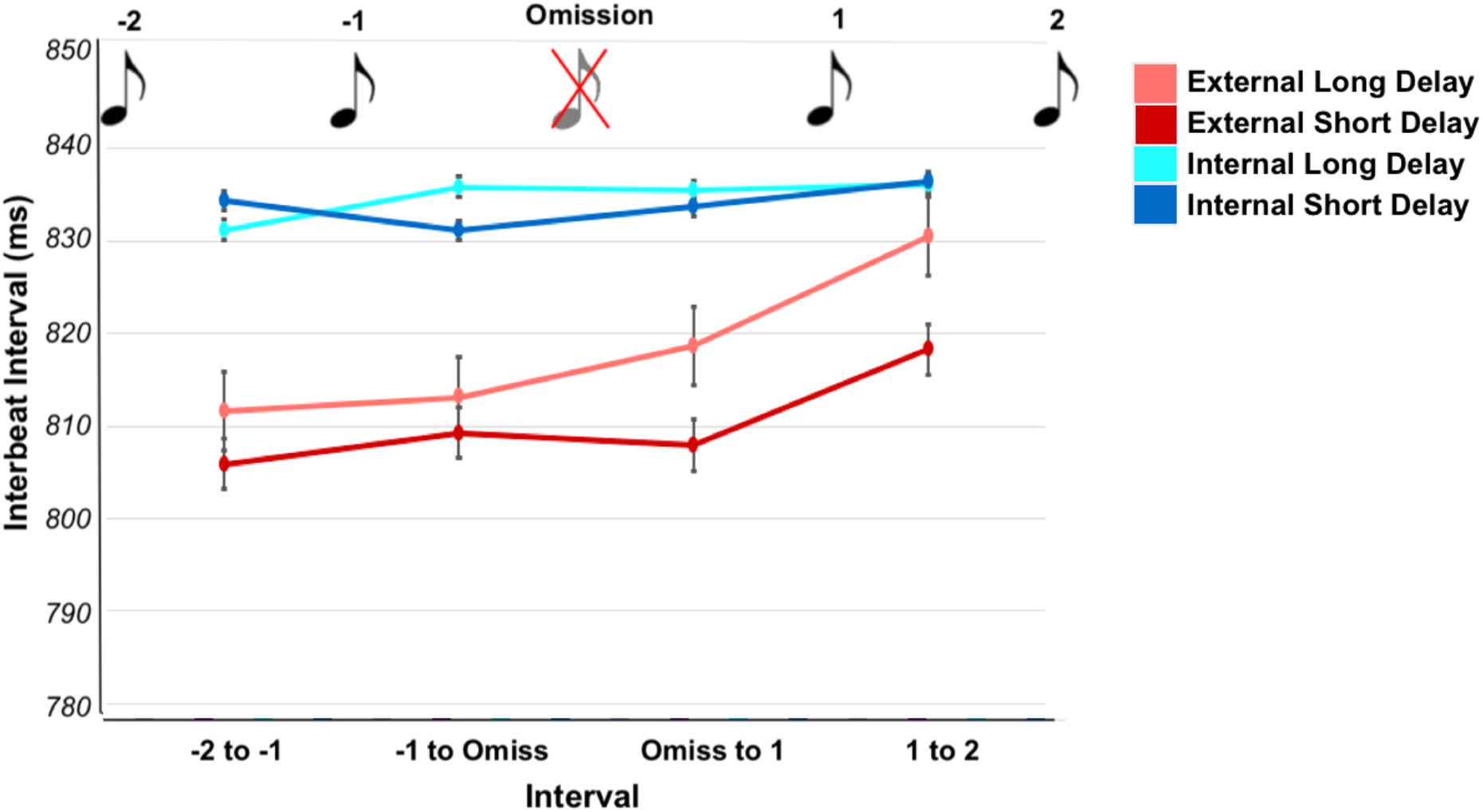
Interbeat intervals in relation to the omission, with error bars reflecting standard error.

Posthoc t-tests revealed that the first IBI after the omission (IBI omission to 1) was significantly faster (short delay: M=818.674, SD=106.089; long delay: M=807.908, SD=96.939) than the following IBI (IBI 1 to 2) (short delay: M=830.529, SD=107.646; long delay: M=818.255, SD=100.182) for external attention trials during both cardio-audio short delay stimulation (t(33) = −4.820, *p* <.001) and long delay stimulation (t(33) = −3.535, *p* = .001). There was no significant difference between the post-omission IBIs during internal attention trials ((short delay: t(33) = −0.981, *p* =.334); (long delay: W = 234, *p* =.285)). This appears to reflect a cardiac deceleration when the omission was a target (i.e. during external attention) (see Figure 5).

## Discussion

Several theories propose that emotion and embodied self-awareness arise from the integration of internal and external signals and their respective precision-weighted expectations (Barrett & Simmons, 2015; Seth, 2013; Seth et al., 2012; Seth & Friston., 2016). Here we investigated these mechanisms of integrated interoceptive and exteroceptive expectations by comparing HEPs during heartbeat-predicted omissions, thus allowing a measure of pure prediction signals without the contamination of bottom-up auditory inputs (Chennu et al., 2016; Wacongne et al., 2011).

First, we observed a pre-omission HEP difference when comparing cardio-audio delay trials, reflected in qualitatively different topographical distributions (see Figure 2A). Consistent with the hypothesis that interoceptive signals guide expectations of exteroceptive stimuli, this result indicates that different expectations of upcoming sounds are induced by different cardio-audio delays, and that these differential expectations are supported by not entirely overlapping regions of cortex. Pfeiffer & De Lucia (2017) reported a similar HEP difference during omission periods when comparing cardio-audio synchronous stimulation with asynchronous stimulation, supporting the integration of cardiac signals to predict auditory stimuli. However, because the sounds in that study (and therefore omission responses) were time-locked to the R-peak during synchronous stimulation but shuffled relative to the R-peak in the asynchronous condition, the auditory omission response is confounded in that contrast. We control for this in our study by comparing trials with sounds at fixed cardio-audio intervals, ensuring the auditory omission response is time-locked to the heartbeat in both delay conditions. This allows for the comparison of pre-omission periods, and later omission-locked responses, which subsequently excludes the auditory omission response as a confound. Nevertheless, our HEP differences across perceived synchrony are consistent with that reported by Pfeiffer & De Lucia (2017). Similarly, in another study consistent with heartbeat-driven auditory predictions, Van Elk et al., (2014) observed a weak auditory N1 suppression to heartbeat-locked sounds, in comparison to cardio-audio asynchronous sounds, although not statistically significant in that study (p = .07).

We also observe an interaction between attention and cardio-audio delay when comparing HEPs locked to the onset of the omission. This is present as a larger positivity to short delay omissions than long delay omissions, when attending internally only. This supports our hypothesis of stronger unfulfilled expectations of a tone in trials presenting sounds at a short perceived synchronous delay than at a longer perceived asynchronous delay. These results are additionally consistent with the role of top-down attentionally-mediated mechanisms in generating expectations of upcoming stimuli. This is supported by modelling evidence, highlighting that omissions are generated by top-down driving inputs, which are attentionally modulated via strengthened downward connections (Chennu et al., 2016). Additionally, attention has been found to enhance mismatch and omission responses, further supporting the role of attention at modulating predictive mechanisms (Chennu et al., 2013; 2016; Garrido et al., 2017; Raij et al., 1997). Despite this, Pfeiffer & De Lucia (2017) reported a heart-beat driven prediction error effect in a group of participants who were naive to the presence of omissions, contrary to our results of absent heartbeat-driven effects when not attending to the heartbeat. Nevertheless, our observation that attention did not modulate the magnitude of our early ERP effect but did modulate the amplitude of the later ERP effect is consistent with the view that early ERPs reflect unconscious/preconscious processing, while later ERPs reflect ignition of representations into consciousness and processing in service of task demands (Dehaene et al., 1998; Sergent et al., 2005).

The modulating nature of attention on HEPs is consistent with previous research and with the interpretation of the HEP as a marker of precision-weighted prediction error of each individual heartbeat (García-Cordero et al., 2017; Montoya et al., 1993; Petzschner et al., 2019; Schandry et al., 1986; Villena-González et al., 2017; Yuan et al., 2007). Attention is proposed to modulate predictive mechanisms by enhancing the precision of attended prediction errors, relative to the precision of their priors (Ainley et al., 2016; Hohwy, 2012; Petzschner et al., 2019). Subsequently, attending to internal signals could enhance the precision of interoceptive prediction errors, resulting in their propagation up the predictive hierarchy to update models for more accurate future predictions regarding each heartbeat. The enhanced cardiac predictions would in turn allow for more precise auditory predictions of heartbeat-locked sounds, such as those presented in our task. The larger positivity to short-delay omissions may be because heartbeat-driven predictions of external stimuli are only stable/accurate across relatively short intervals from the heartbeat (i.e. ~250ms). Similarly, Critchley et al., (2004) found a greater difference in fMRI activity between cardio-audio delay conditions when attending internally, than externally. This was reflected as an increase in the frontal operculum and insula, dorsal and medial parietal lobe, right dorsolateral prefrontal cortex, dorsal cingulate, and lateral temporal cortices during internal attention relative to external. This cortical network overlaps broadly with the source estimates of our interaction of attention with cardio-audio delay in the right inferior frontal gyrus, right anterior prefrontal cortex, bilateral premotor cortex, bilateral primary motor cortex and ventral temporal cortex.

As individual differences in the ability to perceive heartbeat sensations can also be framed as differences in precision, we expected interoceptive accuracy and awareness to similarly modulate interoceptive predictive mechanisms. However, we found no relationship between interoceptive ability and the HEP differences observed in our task, with the exception of a moderate negative correlation of interoceptive accuracy with the delay effect during external attention. However, this apparent correlation should be interpreted with caution as it was part of an exploratory analysis and failed to pass our significance threshold when corrected for multiple comparisons. The lack of evidence for a relationship between our ERP effects and participants’ interoceptive abilities during internal attention is inconsistent with previous evidence that interoceptive accuracy modulated HEP responses (Katkin, Cestaro, & Weitkunat, 1991; Pollatos, Kirsch, & Schandry, 2005; Pollatos & Schandry, 2004; Schandry et al., 1986). However, previous research used heartbeat counting tasks to assess interoceptive performance, rather than the heartbeat discrimination task used in our study, which likely confounds ability to estimate heart rate or time with the ability to sense individual heartbeats (Brener & Ring, 2016; Corneille et al., 2020; Ring & Brener, 2018). The lack of observed differences between interoceptive ability groups in our study could also be because of individual differences in the timing of heartbeat sensations, likely due to biological differences (Wiens & Palmer, 2001). Therefore, some individuals may have performed poorly because they perceived both delay conditions as asynchronous (Brener et al., 1993; Brener & Ring., 2016). This could be investigated in future research by previously determining each individual’s perceived synchronous delay (using the method of constant stimuli (Brener et al., 1993), for example) and subsequently individually adjusting the ‘perceived synchronous’ cardio-audio delay used for each individual (Mesas & Chica., 2003; Brener & Kluvitse., 1988). It’s also possible that HEP differences related to interoceptive ability occur at later latencies than we could measure in our design. For example, ERPs related to metacognition are thought to occur at late latencies (between 550-1900ms) which would overlap with ERPs evoked by successive auditory stimuli in our design (Skavhaug et al., 2010; Sommer et al., 1995; Tsalas et al., 2018).

A potential limitation of our task design is that the internal and external tasks differ in their difficulty. However, we argue that if our observed HEP differences are the result of a task difficulty confound then we would expect that these effects would also correlate with interoceptive performance, which they don’t. A further potential limitation is that the omission is task-relevant in the external task only, perhaps reflected in the post-omission cardiac deceleration during external trials. However, we do not observe any HEP differences as a result of cardio-audio delay during external attention, which would not be expected if task relevance of the omission were an influence on the predictive effects reflected in the HEP. Future research could use an alternative external task of increased difficulty with equal omission task-relevance, such as determining the synchronicity of sounds with a faint flashing visual stimulus, excluding task-related differences as a potential confound.

Previous research has stressed the importance of controlling for ECG artefacts when comparing HEP responses (Kern et al., 2013; Van Elk et al., 2014). We corrected for ECG artefacts using a similar method to that used by Van Elk et al (2014), by subtracting the average HEP response during rest periods for each participant. Our correction was potentially more conservative as it was time locked to each heartbeat within individual trials. Considering the ECG correction applied, and the lack of statistical difference between ECG responses across conditions of interest, we conclude that our observed HEP differences are unlikely to be due to differences in ECG activity, but rather reflect predictive mechanisms of the integration of internal and external stimuli.

Our results support the mechanisms underlying interoceptive predictive coding accounts that suggest that embodied selfhood and emotional experience are a result of integrated self-related predictions from multiple modalities (including interoceptive, exteroceptive and proprioceptive signals) (Barrett & Simmons, 2015; Seth, 2013; Seth, Suzuki, & Critchley, 2012; Seth & Friston., 2016). This is supported by studies which demonstrated the contribution of integrative interoceptive signals with visual cues to enhance body ownership and self-recognition (Aspell et al., 2013; Heydrich et al., 2018; Sel et al., 2017; Suzuki et al., 2013). Additionally, interoceptive and exteroceptive integration has been suggested to explain the generation of a first-person perspective, describing how our unified conscious experience of the external world is integrated with the experience of the self, with particular focus on interoception as a binding agent (Azzalini et al., 2019). These viewpoints, therefore, demonstrate the potential function of the integrated interoceptive and exteroceptive mechanisms observed in our study.

Investigating HEP differences across cardio-audio delay conditions may be a useful clinical tool for assessing dysfunctional interoceptive-exteroceptive predictive mechanisms. As mentioned, the experience of emotion or selfhood is proposed to be the result of the integration of interoceptive predictive mechanisms with exteroception and proprioception (Seth & Friston., 2016). Therefore, measuring pure predictive signals during omissions, which reflect interoceptive and exteroceptive integration, may be useful for diagnosing dissociative disorders, schizophrenia, or anxiety (Paulus & Stein, 2010; Petzschner et al., 2017; Seth, 2013; Seth et al., 2012; Synofzik et al., 2010). Additionally, if interoceptive and exteroceptive integrative mechanisms contribute towards a unified conscious first-person perspective, then observing preserved mechanisms could be useful for diagnosing awareness in patients with disorders of consciousness (Azzalini et al., 2019). This would be advantageous because current methods of assessing awareness focus almost exclusively on responses to external stimuli, whereas assessing interoceptive and exteroceptive integration could provide a method of assessing both external perceptual and internal self-related aspects of awareness.

In conclusion, our results demonstrate that interoceptive signals can guide expectations of exteroceptive stimuli and that attentional-precision modulates integrative cross-modal predictive mechanisms. Nevertheless, we found no evidence that the HEPs were related to subjective experience of heartbeat sensations suggesting low validity of our two-alternative-forced-choice method of assessing interoceptive awareness, or that there exists a more subtle interaction of HEPs and subjective experience. The integrative interoceptive and exteroceptive predictive mechanisms described here provide a useful tool for assessing embodied and interoceptive predictive coding accounts of cognition and clinical disorders.

## Acknowledgments

This work was supported by generous funding from the Medical Research Council IMPACT Doctoral Training Programme at the University of Birmingham (Scholarship to LB) and a Medical Research Council New Investigator Research Grant (MR/P013228/1; to Principal investigator DC).

## Data availability

All data and scripts are available at [https://osf.io/v9khb/].

## References

Adams RA, Shipp S, Friston KJ. 2013. Predictions not commands: Active inference in the motor system. Brain Structure & Function. 218(3):611–643.

Ainley V, Apps MAJ, Fotopoulou A, Tsakiris M. 2016. ‘Bodily precision’: A predictive coding account of individual differences in interoceptive accuracy. Philosophical Transactions of the Royal Society B: Biological Sciences. 371(1708):20160003.

Allen M, Friston KJ. 2018. From cognitivism to autopoiesis: Towards a computational framework for the embodied mind. Synthese. 195(6):2459–2482.

Aspell JE, Heydrich L, Marillier G, Lavanchy T, Herbelin B, Blanke O. 2013. Turning body and self inside out: visualized heartbeats alter bodily self-consciousness and tactile perception. Psychological science. 24(12):2445–2453.

Azzalini D, Rebollo, I, Tallon-Baudry, C. 2019. Visceral Signals Shape Brain Dynamics and Cognition. Trends in Cognitive Sciences. 23(6):488–509.

Barrett LF, Simmons WK. 2015. Interoceptive predictions in the brain. Nature Reviews. Neuroscience. 16(7):419–429.

Brener J, Kluvitse C. 1988. Heartbeat Detection: Judgments of the Simultaneity of External Stimuli and Heartbeats. Psychophysiology. 25(5):554–561.

Brener J, Liu X, Ring, C. 1993. A method of constant stimuli for examining heartbeat detection: Comparison with the Brener-Kluvitse and Whitehead methods. Psychophysiology. 30(6):657–665.

Brener J, Ring, C. 2016. Towards a psychophysics of interoceptive processes: the measurement of heartbeat detection. Philosophical Transactions of the Royal Society B: Biological Sciences. 371(1708):20160015.

Cameron OG. 2001. Interoception: The inside story–a model for psychosomatic processes. Psychosomatic Medicine. 63(5):697–710.

Chennu S, Noreika V, Gueorguiev D, Shtyrov Y, Bekinschtein TA, Henson R. 2016. Silent Expectations: Dynamic Causal Modeling of Cortical Prediction and Attention to Sounds That Weren’t. Journal of Neuroscience, 36(32). 8305–8316.

Corneille O, Desmedt O, Zamariola G, Luminet O, Maurage P. 2020. A heartfelt response to Zimprich et al.(2020), and Ainley et al.(2020)’s commentaries: Acknowledging issues with the HCT would benefit interoception research. Biological Psychology. 152:107869.

Craig, ADB. 2009. How do you feel--now? The anterior insula and human awareness. Nature Reviews. Neuroscience. 10(1):59–70.

Critchley HD, Harrison NA. 2013. Visceral Influences on Brain and Behavior. Neuron, 77(4):624–638.

Critchley HD, Wiens S, Rotshtein P, Öhman A, Dolan RJ. 2004. Neural systems supporting interoceptive awareness. Nature Neuroscience. 7(2):189–195.

Damasio A, Carvalho GB. 2013. The nature of feelings: Evolutionary and neurobiological origins. Nature Reviews Neuroscience. 14(2):143–152.

Dehaene S, Naccache L, Le Clec’H G, Koechlin E, Mueller M, Dehaene-Lambertz G, … Le Bihan D. 1998. Imaging unconscious semantic priming. Nature. 395(6702):597–600.

Ewing DL, Manassei M, van Praag CG, Philippides AO, Critchley HD, Garfinkel SN. 2017. Sleep and the heart: Interoceptive differences linked to poor experiential sleep quality in anxiety and depression. Biological psychology. 127:163–172.

Faul F, Erdfelder E, Lang AG, Buchner A. 2007. G* Power 3: A flexible statistical power analysis program for the social, behavioral, and biomedical sciences. Behavior research methods. 39(2):175–191.

Feldman, H, Friston KJ. 2010. Attention, Uncertainty, and Free-Energy. Frontiers in Human Neuroscience. 4.

Forkmann T, Scherer A, Meessen J, Michal M, Schächinger H, Vögele C, Schulz A. 2016. Making sense of what you sense: Disentangling interoceptive awareness, sensibility and accuracy. International Journal of Psychophysiology. 109:71–80.

Friston K. (2009). The free-energy principle: A rough guide to the brain? Trends in Cognitive Sciences. 13(7):293–301.

Friston K. 2010. The free-energy principle: A unified brain theory? Nature Reviews Neuroscience. 11(2): 127–138.

García-Cordero I, Esteves S, Mikulan E, Hesse E, Baglivo FH, Silva W, … Sedeño L. 2017. Attention, in and Out: Scalp-Level and Intracranial EEG Correlates of Interoception and Exteroception. Frontiers in neuroscience. 11:411.

Garfinkel SN, Seth AK, Barrett AB, Suzuki K, Critchley HD. 2015. Knowing your own heart: distinguishing interoceptive accuracy from interoceptive awareness. Biological psychology. 104:65–74.

Garrido MI, Kilner JM, Stephan KE, Friston KJ. 2009. The mismatch negativity: a review of underlying mechanisms. Clinical neurophysiology. 120(3):453–463.

Gray MA, Harrison NA, Wiens S, Critchley HD. 2007. Modulation of Emotional Appraisal by False Physiological Feedback during fMRI. PLoS ONE. 2:145–172.

Haker H, Schneebeli M, Stephan KE. 2016. Can Bayesian Theories of Autism Spectrum Disorder Help Improve Clinical Practice? Frontiers in Psychiatry. 7.

Henson RN, Mattout J, Phillips C, Friston KJ. 2009. Selecting forward models for MEG source-reconstruction using model-evidence. Neuroimage. 46(1): 168–176.

Heydrich L, Aspel JE, Marillier G, Lavanchy T, Herbelin B, Blanke O. 2018. Cardio-visual full body illusion alters bodily self-consciousness and tactile processing in somatosensory cortex. Scientific reports. 8(1): 1–8.

Hohwy J. 2012. Attention and Conscious Perception in the Hypothesis Testing Brain. Frontiers in Psychology. 3.

Katkin ES, Cestaro VL, Weitkunat R. 1991. Individual differences in cortical evoked potentials as a function of heartbeat detection ability. The International Journal of Neuroscience. 61(3-4): 269–276.

Kern M, Aertsen A, Schulze-Bonhage A, Ball T. 2013. Heart cycle-related effects on event-related potentials, spectral power changes, and connectivity patterns in the human ECoG. Neuroimage, 81:178–190.

Kligfield P, Gettes LS, Bailey JJ, Childers R, Deal BJ, Hancock EW, … Wagner GS. 2007. Recommendations for the Standardization and Interpretation of the Electrocardiogram. Journal of the American College of Cardiology. 49(10):1109–1127.

Kothe C, Medine D, Grivich M. 2018. Lab Streaming Layer 2014. URL: https://github.com/sccn/labstreaminglayer (visited on 26/02/2020).

Lawson RP, Rees G, Friston KJ. 2014. An aberrant precision account of autism. Frontiers in Human Neuroscience, 8.

López JD, Litvak V, Espinosa JJ, Friston K, Barnes GR. 2014. Algorithmic procedures for Bayesian MEG/EEG source reconstruction in SPM. NeuroImage, 84:476–487.

Macmillan NA, Creelman CD. 1990. Response bias: Characteristics of detection theory, threshold theory, and nonparametric indexes. Psychological Bulletin. 107(3):401.

Maniscalco B, Lau H. 2012. A signal detection theoretic approach for estimating metacognitive sensitivity from confidence ratings. Consciousness and Cognition. 21(1):422–430.

Maniscalco B, Lau H. 2014. Signal Detection Theory Analysis of Type 1 and Type 2 Data: Meta-d’, Response-Specific Meta-d’, and the Unequal Variance SDT Model. In: Fleming SM, Frith CD. The Cognitive Neuroscience of Metacognition. Springer, Berlin, Heidelberg. p 25–66.

Maris E, Oostenveld R. 2007. Nonparametric statistical testing of EEG-and MEG-data. Journal of Neuroscience Methods. 164(1): 177–190.

Mesas AA, Chica JP. 2003. Facilitation of heartbeat self-perception in a discrimination task with individual adjustment of the S+ delay values. Biological psychology. 65(1):67–79.

Mognon A, Jovicich J, Bruzzone L, Buiatti M. 2011. ADJUST: An automatic EEG artifact detector based on the joint use of spatial and temporal features: Automatic spatio-temporal EEG artifact detection. Psychophysiology. 48(2):229–240.

Montoya P, Schandry R, Müller A. 1993. Heartbeat evoked potentials (HEP): Topography and influence of cardiac awareness and focus of attention. Electroencephalography and Clinical Neurophysiology/Evoked Potentials Section. 88(3):163–172.

Nakamura M, Shibasaki H. 1987. Elimination of EKG artifacts from EEG records: a new method of non-cephalic referential EEG recording. Electroencephalography and clinical neurophysiology. 66(1):89–92.

Nolan H, Whelan R, Reilly RB. 2010. FASTER: Fully Automated Statistical Thresholding for EEG artifact Rejection. Journal of Neuroscience Methods. 192(1): 152–162.

Oostenveld R, Fries P, Maris E, Schoffelen JM. 2011. FieldTrip: Open Source Software for Advanced Analysis of MEG, EEG, and Invasive Electrophysiological Data. Computational Intelligence and Neuroscience. 2011:1–9.

Paulus MP, Stein MB. 2010. Interoception in anxiety and depression. Brain structure and Function. 214(5-6):451–463.

Petzschner FH, Weber LA, Gard T, Stephan KE. 2017. Computational psychosomatics and computational psychiatry: toward a joint framework for differential diagnosis. Biological Psychiatry. 82(6):421–430.

Petzschner FH, Weber LA, Wellstein KV, Paolini G, Do CT, Stephan KE. 2019. Focus of attention modulates the heartbeat evoked potential. NeuroImage. 186:595–606.

Pfeiffer C, De Lucia M. 2017. Cardio-audio synchronization drives neural surprise response. Scientific reports. 7(1):1–10.

Pollatos O, Kirsch W, Schandry R. 2005. Brain structures involved in interoceptive awareness and cardioafferent signal processing: A dipole source localization study. Human Brain Mapping. 26(1):54–64.

Pollatos O, Schandry R. 2004. Accuracy of heartbeat perception is reflected in the amplitude of the heartbeat-evoked brain potential: Heartbeat-evoked potential and heartbeat perception. Psychophysiology. 41(3):476–482.

Porges S. 1993. Body perception questionnaire. Laboratory of Developmental Assessment, University of Maryland. Pridobljeno. 15(12):2009.

Price D. 2020. MNI2FS: High Resolution Surface Rendering of MNI Registered Volumes (https://www.github.com/dprice80/mni2fs). GitHub. Retrieved April 3, 2020.

Quadt L, Critchley HD, Garfinkel SN. 2018. The neurobiology of interoception in health and disease: Neuroscience of interoception. Annals of the New York Academy of Sciences. 1428(1):112–128.

Quattrocki E, Friston K. 2014. Autism, oxytocin and interoception. Neuroscience & Biobehavioral Reviews. 47:410–430.

Raimondo F, Rohaut B, Demertzi A, Valente M, Engemann DA, Salti M, … Sitt JD. 2017. Brain–heart interactions reveal consciousness in noncommunicating patients. Annals of neurology. 82(4):578–591.

Raij T, McEvoy L, Mäkelä JP, Hari R. 1997. Human auditory cortex is activated by omissions of auditory stimuli. Brain research. 745(1-2):134–143.

Rao RPN, Ballard DH. 1999. Predictive coding in the visual cortex: A functional interpretation of some extra-classical receptive-field effects. Nature Neuroscience. 2(1):79–87.

Ring C, Brener J. 2018. Heartbeat counting is unrelated to heartbeat detection: A comparison of methods to quantify interoception. Psychophysiology, 55(9):13084.

Salomon R, Ronchi R, Dönz J, Bello-Ruiz J, Herbelin B, Martet R, … Blanke O. 2016. The Insula Mediates Access to Awareness of Visual Stimuli Presented Synchronously to the Heartbeat. The Journal of Neuroscience. 36(18):5115–5127.

Schandry R, Sparrer B, Weitkunat R. 1986. From the heart to the brain: A study of heartbeat contingent scalp potentials. International Journal of Neuroscience. 30(4):261–275.

Sel A, Azevedo RT, Tsakiris M. 2017. Heartfelt self: cardio-visual integration affects self-face recognition and interoceptive cortical processing. Cerebral Cortex. 27(11):5144–5155.

Sergent C, Baillet S, Dehaene S. 2005. Timing of the brain events underlying access to consciousness during the attentional blink. Nature neuroscience. 8(10):1391–1400.

Seth AK. 2013. Interoceptive inference, emotion, and the embodied self. Trends in Cognitive Sciences, 17(11):565–573.

Seth AK, Friston KJ. 2016. Active interoceptive inference and the emotional brain. Philosophical Transactions of the Royal Society B: Biological Sciences, 371(1708).

Seth AK, Suzuki K, Critchley HD. 2012. An Interoceptive Predictive Coding Model of Conscious Presence. Frontiers in Psychology. 2.

Sommer W, Heinz A, Leuthold H, Matt J, Schweinberger SR. 1995. Metamemory, distinctiveness, and event-related potentials in recognition memory for faces. Memory & Cognition. 23(1):1–11.

Sherrington C. 1952. The integrative action of the nervous system. CUP Archive.

Skavhaug IM, Wilding EL, Donaldson DI. 2010. Judgments of learning do not reduce to memory encoding operations: Event-related potential evidence for distinct metacognitive processes. Brain research. 1318:87–95.

Suzuki K, Garfinkel SN, Critchley HD, Seth AK. 2013. Multisensory integration across exteroceptive and interoceptive domains modulates self-experience in the rubber-hand illusion. Neuropsychologia. 51(13):2909–2917.

Synofzik M, Thier P, Leube DT, Schlotterbeck P, Lindner A. 2010. Misattributions of agency in schizophrenia are based on imprecise predictions about the sensory consequences of one’s actions. Brain. 133(1):262–271.

Tsakiris M, Critchley H. 2016. Interoception beyond homeostasis: Affect, cognition and mental health. Philosophical Transactions of the Royal Society B: Biological Sciences, 371(1708):20160002.

Tsalas NR, Müller BC, Meinhardt J, Proust J, Paulus M, Sodian B. 2018. An ERP study on metacognitive monitoring processes in children. Brain research. 1695:84–90.

Van Elk M, Lenggenhager B, Heydrich L, Blanke O. 2014. Suppression of the auditory N1-component for heartbeat-related sounds reflects interoceptive predictive coding. Biological psychology. 99:172–182.

Villena-González M, Moënne-Loccoz C, Lagos RA, Alliende LM, Billeke P, Aboitiz F, … Cosmelli D. 2017. Attending to the heart is associated with posterior alpha band increase and a reduction in sensitivity to concurrent visual stimuli. Psychophysiology/ 54(10):1483–1497.

Wacongne C, Labyt, E, van Wassenhove V, Bekinschtein T, Naccache L, Dehaene S. 2011. Evidence for a hierarchy of predictions and prediction errors in human cortex. Proceedings of the National Academy of Sciences. 108(51):20754–20759.

Wiens S, Palmer SN. 2001. Quadratic trend analysis and heartbeat detection. Biological Psychology. 58:159–175.

Yuan H, Yan HM, Xu XG, Han F, Yan Q. 2007. Effect of heartbeat perception on heartbeat evoked potential waves. Neuroscience Bulletin. 23(6):357–362.

